# AIRE is induced in oral squamous cell carcinoma and promotes cancer gene expression

**DOI:** 10.1101/760959

**Authors:** Chi Thi Kim Nguyen, Wanlada Sawangarun, Masita Mandasari, Kei-ichi Morita, Kou Kayamori, Akira Yamaguchi, Kei Sakamoto

**Affiliations:** Department of Oral Pathology, Graduate School of Medical and Dental Sciences, Tokyo Medical and Dental University, Tokyo, Japan; Department of Maxillofacial Surgery, Graduate School of Medical and Dental Sciences, Tokyo Medical and Dental University, Tokyo, Japan; Bioresource Research Center, Tokyo Medical and Dental University, Tokyo, Japan; Oral Health Science Center, Tokyo Dental College, Tokyo, Japan

**Author notes:** Corresponding author, Kei Sakamoto, Department of Oral Pathology, Graduate School of Medical and Dental Sciences, Tokyo Medical and Dental University, 1-5-45 Yushima, Bunkyo-ku, Tokyo 113-0034, Japan.

## Abstract

Autoimmune regulator (AIRE) is a transcriptional regulator that is primarily expressed in medullary epithelial cells, where it induces tissue-specific antigen expression. Under pathological conditions, AIRE expression is induced in epidermal cells and promotes skin tumor development in association with stress-responsive keratin KRT17. This study aimed to clarify the role of AIRE in the pathogenesis of oral squamous cell carcinoma (OSCC). AIRE expression was evaluated in seven OSCC cell lines and in OSCC tissue specimens. Transient or constitutive expression of AIRE in 293A cells induced KRT17 expression. cDNA microarray analysis of 293A cells stably expressing AIRE revealed that STAT1 and ICAM1 were significantly upregulated by AIRE. Expression of KRT17, STAT1, ICAM1, MMP9, CXCL10, and CXCL11 was elevated in 293A cells stably expressing AIRE, and conversely, was decreased in AIRE-knockout HSC3 OSCC cells when compared to the respective controls. Upregulation of KRT17, STAT1, and ICAM in OSCC cells was confirmed in tissue specimens by immunohistochemistry. We provide evidence that AIRE exerts transcriptional control in cooperation with ETS1. Expression of STAT1, ICAM1, CXCL10, and MMP9 was increased in 293A cells upon *Ets1* transfection, and coexpression of AIRE resulted in enhanced expression of STAT1. AIRE coprecipitated with ETS1 in a modified immunoprecipitation assay using formaldehyde crosslinking. Chromatin immunoprecipitation and quantitative PCR analysis revealed that promoter fragments of STAT1, ICAM1, CXCL10, and MMP9 were enriched in the AIRE precipitates. In oral cancer cells, ETS1 was diffusely located in the nucleus and partially overlapped with dot-like AIRE accumulation sites. Nuclear translocation of AIRE was promoted by cotransfection with *Ets1*. These results indicate that AIRE is induced in OSCC and supports cancer-related gene expression in cooperation with ETS1. This is a novel function of AIRE in extrathymic tissues under the pathological condition.

## Introduction

The burden of oral cancer has significantly increased in many parts of the world, causing more than 145.000 death in 2012 worldwide [1]. Despite the advances in modern medicine, oral cancer can have devastating effects on critical life functions. About 90 percent of oral cancers are squamous cell carcinomas (OSCCs) that arise from keratinocytes of the oral mucosa.

Cancer development requires the acquisition of several properties, such as unlimited proliferation, vascularization, and invasion [2]. Aberrant growth control in cancer cells is the consequence of accumulated disorders in multiple cell-regulatory systems. Molecular analysis of OSCCs has uncovered genetic and epigenetic alterations in several distinct driver pathways, including mitogenic signaling and cell-cycle regulatory pathways, which lead to continual and unregulated proliferation of cancer cells. In addition, cancer cells recruit surrounding non-transformed cells, such as stromal cells and inflammatory cells, to create a tumor microenvironment that fosters tumor growth and invasion. Under the influence of interaction with the tumor microenvironment, cancer cells express a unique group of proteins that are absent or expressed at very low levels in normal cells. Differentially expressed proteins in OSCC compared to normal keratinocytes include various cytokines, chemokines (for example, CXCLs), extracellular matrix proteins, matrix remodeling enzymes (for example, matrix metalloproteases (MMPs)), cell adhesion molecules, and cytoskeletal proteins [3, 4]. Among these factors that characterize cancer cells, we have been interested in keratin 17 (KRT17) because it is consistently and strongly induced in OSCC, and because of its unique functions. Keratins are a family of intermediate filaments that are indispensable for the structural stability of epithelial cells. KRT17 is regarded a “stress-response keratin” that is not expressed or at a very low level in normal squamous epithelium and is induced under pathological conditions, such as wound healing, inflammation, and cancer. KRT17 exerts diverse functions, including the regulation of Th1-type chemokine expression, which promotes tumor growth [5]. Interestingly, KRT17 has been recently reported to interact with AIRE [6]. *AIRE* was first identified as a causative gene for autoimmune polyendocrinopathy-candidiasis-ectodermal dystrophy syndrome, which gives rise to multiple endocrine disorders, chronic mucocutaneous candidiasis, and various ectodermal defects.

AIRE consists of four domains, including a caspase recruitment domain/homogeneously staining region (CARD/HSR) domain that drives nuclear localization and oligomerization, an Sp100, AIRE, NucP41/75, DEAF1 (SAND) domain, and plant homeodomain 1 (PHD1) and PHD2 domains. The closest homolog of AIRE is Sp100, which shares the CARD, SAND, and PHD domains [7]. AIRE localizes in the nucleus in discrete, punctate structures that resemble promyelocytic leukemia nuclear bodies, which contain Sp100 family members. The nuclear localization and the presence of domains that are shared by transcription factors suggested a role for AIRE in the regulation of gene expression, and indeed, numerous *in-vitro* studies have demonstrated that AIRE functions as a transcriptional activator [7]. AIRE is expressed in medullary thymic epithelial cells (mTECs), in which it induces ectopic expression of peripheral tissue-specific antigens, thereby facilitating the elimination of self-reactive T cells. AIRE expression has been also reported in extrathymic tissues, including epidermal keratinocytes [8–11]. In the mouse epidermis, Aire is induced by inflammatory or tumorigenic stimuli concurrently with Krt17 and stimulates the transcription of several chemokine genes, such as *Cxcl10* and *Cxcl11*, and a gene encoding matrix-degrading enzyme, *Mmp9*, in keratinocytes [6]. The onset of genetically induced skin tumor is delayed in *Aire*-knockout mice, which mimic the phenotype of *Krt17*-knockout mice [6]. These results suggest a novel role of AIRE in extrathymic epithelial tissues under pathological conditions; however, the significance of AIRE in other cancers remains to be elucidated.

The above findings inspired us to evaluate whether AIRE participates in the pathogenesis of OSCC. To this end, we first assessed the expression of AIRE in OSCC. Then, we explored the role of AIRE in the regulation of *KRT17* and other genes associated with OSCC development.

## Materials and Methods

### Chemically induced mouse model of tongue and esophageal cancer

CD-1 mice were obtained from Sankyo Labo Service Co. (Tokyo, Japan) and were housed in groups of less than five in plastic cages in a temperature-controlled room with 12 h light-12 h dark cycles, and were provided free access to tap water and food. To establish the cancer model, 4-nitroquinoline-1-oxide (4-NQO) was added to the drinking water at a concentration of 100 μg/ml and administered to 6-week-old mice for 8 weeks. After the administration period, normal drinking water was given. The health of the animals was monitored daily. Behavioral changes were used as criteria to determine a humane endpoint. None of the mice reached the endpoint before the day of sacrifice. Mice were euthanized by cervical dislocation at 30 weeks of age, and tongues and esophagi were harvested. The animal studies were reviewed and approved by the institutional animal care and use committee of Tokyo Medical and Dental University (registration numbers: 0150213A and 0160316A).

### Histology and immunohistochemistry

The tissues were fixed in 10% neutral buffered formalin for 24–48 h and embedded in paraffin. Four micrometer-thick tissue sections were cut and stained with hematoxylin and eosin. For immunohistochemistry, the tissue sections were incubated in antigen retrieval buffer (10 mM Tris (pH 9.0), 1 mM EDTA) at 120°C for 15 min. Endogenous peroxidase activity was quenched by 3% H_2_O_2_ for 10 min. The sections were incubated with primary antibody diluted 1:500 in TBST (10 mM Tris (pH 7.4), 150 mM NaCl, 0.1% Tween 20) at 4°C overnight. After washing, the sections were incubated with a secondary antibody at room temperature for 1 h. 3,3′-Diaminobenzidine (DAB) was used for chromogenic detection. Primary antibodies used in this study were anti-AIRE (C-2, Santa Cruz Biotechnology, Santa Cruz, CA, USA) for detection of human AIRE, anti-AIRE (M300, Santa Cruz Biotechnology) for detection of mouse AIRE, anti-keratin 17 (EPR1624Y, Abcam, Cambridge, UK), anti-keratin 17 (polyclonal, #1900, Abcam (Epitomics)), anti-GAPDH (D16H11, Cell Signaling Technology, Danvers, MA, USA), anti-STAT1 alpha (EPYR2154, Abcam), anti-STAT1 phospho S727 (EPR3146, Abcam), anti-ICAM1 (EP1442Y, Abcam), anti-ETS1 (D8O8A, Cell Signaling Technology), anti-keratin 14 (LL002, Abcam), anti-Flag M2 (Sigma-Aldrich, St. Louis, MO, USA), and anti-HA (12CA5, Roche Diagnostics, Basel, Switzerland). Secondary antibodies used in this study were Envision/HRP (Dako, Glostrup, Denmark), HRP-donkey anti-rabbit IgG (Thermo Fisher Scientific, Waltham, MA, USA), and HRP-rabbit anti-mouse IgG, Alexa Fluor 488 goat anti-rabbit IgG, Alexa Fluor 488 goat anti-mouse IgG, Alexa Fluor 594 goat anti-rabbit IgG, and Alexa Fluor 594 goat anti-mouse IgG (Thermo Fisher Scientific).

### Cell culture

Ca9-22 (derived from OSCC of the gingiva) and HSC5 (derived from OSCC of the tongue) cell lines were obtained from the RIKEN Bioresource Center (Tsukuba, Japan). HSC3 (derived from OSCC of the tongue) and HO1N1 (derived from OSCC of the buccal mucosa) were obtained from the Japanese Collection of Research Bioresources (Osaka, Japan). BHY (derived from OSCC of the gingiva) was provided by Dr. Masato Okamoto (Tsurumi Univ.). SAS (derived from OSCC of the tongue) and HSC4 (derived from OSCC of the tongue) were provided from Dr. Masao Saito (Yamanashi Univ.). 293A was purchased from Thermo Fisher Scientific Co. These cells were maintained in Dulbecco’s modified Eagle’s medium: nutrient mixture F-12 (Sigma-Aldrich) with 10% fetal bovine serum. Primary human keratinocytes isolated from neonatal foreskins were obtained from Kurabo Co. (Osaka, Japan) and were maintained in HuMedia-KG2 (Kurabo).

### Plasmid constructs

A pET32a plasmid containing human *AIRE* cDNA was provided by Dr. Yoshitaka Yamaguchi (Keio Univ). The open reading frame of *AIRE* was recovered by PCR using this plasmid as a template and PrimeSTAR GXL DNA Polymerase (Takara, Shiga, Japan). The PCR product was digested with *Eco*R1 and inserted into pcDNA6-3xFlagN (in-house plasmid made using pcDNA6/V5-HisA (Thermo Fisher Scientific) as a backbone vector), or pAcGFP-C2 (Takara) to create N-terminally 3xFlag-tagged AIRE (*Flag-AIRE*), and GFP-tagged AIRE (*GFP-AIRE*), respectively. The plasmid construct targeting exon 2 of human *AIRE* (*AIREKO*) was created using pSpCas9(BB)-2A-GFP (a gift from Dr. Feng Zhang, #48138, Addgene, http://n2t.net/addgene:48138 ; RRID:Addgene_48138) as a backbone vector [12]. The guide sequence was GGAGCGCTATGGCCGGCTGC. pCMV-mFlagEts1 was a gift from Dr. Barbara Graves (# 86099, Addgene; http://n2t.net/addgene:86099; RRID:Addgene_86099) [13]. To remove the Flag tag, a fragment containing the coding region of mouse *Ets1* was amplified by PCR, digested with *Bam*H1 and *Spe*1, and inserted into pcDNA6/V5-HisA.

### Generation of AIRE-overexpressing and AIRE-knockout cells

Transfection was carried out using Polyethylenimine Max (Polysciences, Warrington, PA, USA). To establish AIRE-overexpressing clones, 293A cells were transfected with *Flag-AIRE* and selected on blasticidin (5 µg/ml), and clones were isolated from colonies. To establish knockout clones, HSC3 cells were transfected with *AIREKO*, GFP-positive cells were sorted using BD FACSAria II (BD Biosciences, San Jose, CA, USA) 72 h after transfection to compensate for the low transfection efficiency of HSC3 cells, and expanded through limiting dilution. Clones harboring a nonsense mutation were determined by PCR-direct sequencing using BigDye Terminator v3.1 Cycle Sequencing Kit (Thermo Fisher Scientific).

### Immunofluorescence staining of cultured cells

Cells cultured in a chamber slide were fixed with methanol for 10 min. The cells were permeabilized with 0.5% Triton X-100 for 15 min, washed in PBS, incubated with Image-iT FX Signal Enhancer (Thermo Fisher Scientific) at room temperature for 30 min, and then incubated with primary antibodies (1:500 dilution) for 3 h, followed by incubation with secondary antibodies (1:500 dilution) for 1 h. Nuclei were counterstained with propidium iodide or 4′,6-diamidino-2-phenylindole (DAPI).

### Proliferation and migration assays

Cell proliferation was assessed by measuring metabolic activity using Cell Counting Kit-8 (Dojindo Laboratories, Kumamoto, Japan) according to the manufacturer’s instructions. Transwell migration assays were conducted using ThinCert cell culture inserts with 8 µm pore diameter (Greiner Bio-One, Wemmel, Belgium) according to the manufacturer’s instructions.

### Northern blotting, RT-PCR, real-time RT-PCR, and cDNA microarray analysis

Total RNA was isolated from Ca9-22 and HSC3 cells using NucleoSpin (Macherey-Nagel, Düren, Germany). Northern blot analysis was conducted using DIG Easy Hyb and an RNA probe made with Dig RNA labeling Mixture (Roche Diagnostics, Basel, Switzerland) according to the manufacturer’s protocol. RNA was reverse-transcribed to cDNA using oligo(dT) primers and M-MuLV reverse transcriptase (Thermo Fisher Scientific). PCRs were run with PrimeSTAR GXL DNA Polymerase (Takara) using the following thermal cycling program: 98°C for 1 min, 30 cycles of 98°C for 10 s, 58°C for 15 s, and 68°C for 2 min (*KRT17*) or 20 s (others), and 68°C for 2 min. *GAPDH* was used as an internal control. Quantitative RT-PCR was conducted using Platinum SYBR Green qPCR SuperMix-UDG (Thermo Fisher Scientific) and a LightCycler Nano system (Roche Life Science) using the following thermal cycling program: 95°C for 10 min, 40 cycles of 95°C for 10 s, 60°C for 10 s, and 72°C for 15 s. Primer sequences are available in S1 Table. For cDNA expression microarray analysis using SurePrint G3 Human GE 8×60K Ver.2.0 (Agilent Technologies, Santa Clara, CA, USA), total RNA was sent to a commercial service (Hokkaido System Science, Sapporo, Japan).

### Western blot analysis

Cells were lysed in RIPA buffer (20 mM Tris pH 7.4, 150 mM NaCl, 1% NP40, 0.1% SDS and 0.5% sodium deoxycholate, containing protease inhibitor cocktail cOmplete (Sigma-Aldrich). Protein samples were subjected to 10% SDS-PAGE and transferred to Hybond-ECL (GE Healthcare, Pittsburgh, PA, USA). The blotted membranes were blocked in 2% non-fat milk for 30 min and then incubated with primary antibodies (1:2,000) at 4°C overnight, followed by incubation with secondary antibodies (1:20,000) at room temperature for 1 h, and visualized using ECL Select (GE Healthcare).

### OSCC tissue specimens and immunohistochemistry

The study was reviewed and approved by the institutional ethics committee (registration number: D2012-078). As we used archival tissue specimens obtained for pathology diagnosis, the ethics committee approved waiver of the informed consent requirement in accordance with Ethical Guidelines for Clinical Studies by the Ministry of Health, Labor and Welfare of Japan. OSCC tissue specimens were randomly collected from sixty lateral tongue cancer cases from the archives of the Dental Hospital of Tokyo Medical and Dental University. Three tissue microarrays harboring 20 spots of 4 mm in diameter were constructed as previously described [14]. Fifty one spots were fully evaluable after the loss of spots due to multiple sectioning. Histological evaluation was done as previously described [15]. Cancer-associated inflammation was scored based on the density of inflammatory cell infiltrates around tumor nests at the invasion front. Expression of AIRE, KRT17, ICAM1, pSTAT1 and STAT1 in cancer was scored based on staining intensity in cancer versus normal epithelium, with “+/–” indicating that expression was undetected or detected but without noticeable upregulation in cancer, “+” indicating slight upregulation in cancer as revealed by a noticeable increase in staining intensity in cancer cells, and “++” indicating distinctive upregulation in cancer. All scorings were performed by two pathologists in a blinded manner, and the judgment was discussed until consent was obtained.

### Immunoprecipitation and chromatin immunoprecipitation (ChIP)

Cells were crosslinked with 1% formaldehyde at room temperature for 10 min. The reaction was stopped by incubating the cells with 10% glycine for 15 min. The cells were rinsed twice with PBS and lysed in RIPA buffer. The cell lysate was sonicated for 5 min and centrifuged to pellet the debris. The cleared lysate was divided into two parts, one of which was used for immunoprecipitation and western blot analysis, and the other for immunoprecipitation and DNA quantitative PCR assay. For immunoprecipitation and western blotting, the lysate was incubated with anti-ETS1 antibody (1:100 dilution) at 4°C overnight. Dynabeads Protein G (Thermo Fisher Scientific) were added to the sample, which was incubated for an additional 1 h. The beads were washed with RIPA buffer three times using a magnet stand. Protein was eluted and crosslinks were reversed by heating the beads in SDS gel loading buffer (50 mM Tris-HCl pH6.8, 2% SDS, 10% glycerol, 5% beta-mercaptoethanol, 0.01% bromophenol blue) at 95°C for 10 min. Eluted proteins were subjected to western blot analysis. For ChIP, the lysate was incubated with anti-AIRE antibody at 4°C overnight, then with Dynabeads Protein G for an additional 1 h. The sample was washed first with low-salt wash buffer (20 mM Tris-HCl pH8.0, 150 mM NaCl, 2 mM EDTA, 1% Triton X-100, 0.1% SDS), then with high-salt wash buffer (20 mM Tris-HCl pH8.0, 500 mM NaCl, 2 mM EDTA, 1% Triton X-100, 0.1% SDS), and then with LiCl wash buffer (10 mM Tris-HCl pH8.0, 0.25 M LiCl, 1 mM EDTA, 1% NP-40, 1% sodium deoxycholate). DNA was eluted in elution buffer (100 mM NaHCO_3_, 1% SDS). Crosslinks were reversed by incubation at 65°C overnight, in the presence of RNaseA. Protein in the sample was digested with proteinase K at 60°C for 1 h, and DNA was purified by phenol/chloroform extraction. Primers for quantitative PCR analysis are available in S1 Table.

### Statistical analysis

Quantitative data are presented as the mean ± standard error of at least three replicated experiments. Means were compared using Student’s *t*-test or Mann-Whitney U test. For analysis of relative protein or gene expression data, the ratio t-test was used. *P* < 0.05 was considered statistically significant.

## Results

### AIRE is induced in chemically induced cancer of the upper digestive tract in mice

We first examined AIRE expression in the healthy mouse thymus by immunohistochemical staining. AIRE-expressing cells were detected specifically in some, but not all epithelioid cells in the medulla (Fig 1A). AIRE-positive cells were also positive for keratin 14 (KRT14), confirming that they were mTECs. AIRE localized in the nuclei in punctate regions, which were more clearly observed by fluorescence than by chromogenic detection (Fig 1A). This AIRE protein distribution was in agreement with previous reports [16, 17], validating the immunohistochemical method for AIRE detection.

**Fig 1.**
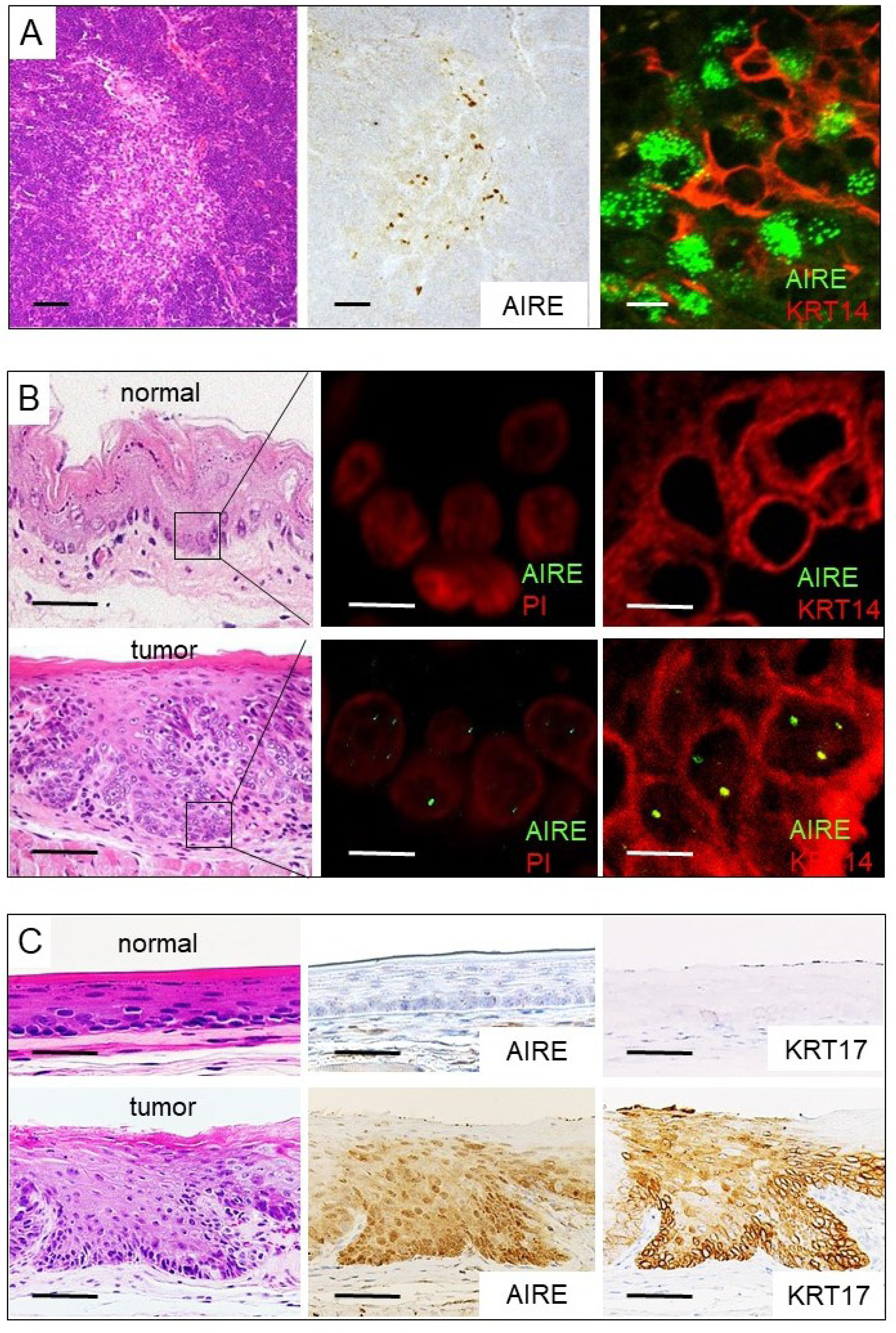
AIRE expression in esophageal tumors in mice. A) AIRE expression in the thymic medullary epithelium. Hematoxylin and eosin staining (left panel), immunohistochemical staining using 3,3’-DAB (middle panel), and immunofluorescence double staining with KRT14 (right panel). Scale bar: 50 µm (left and middle panels), 10 µm (right panel). B) AIRE expression in neoplastic cells of the esophagus. Immunofluorescence staining (middle panel) and immunofluorescence double staining with KRT14 (right panel). PI: propidium iodide. Scale bar: 50 µm (left), 5 µm (middle and right panels). C) Expression of AIRE and KRT17 in neoplastic cells of the esophagus as revealed by immunohistochemical staining using DAB. Scale bar: 50 µm.

We induced tumorigenesis in the upper digestive tract by administering the carcinogen 4-NQO to mice via the drinking water. Multiple tumors developed in the esophagi of all 4-NQO-treated mice (n = 5), whereas no tumor developed in control mice (n = 5). The tumors were histologically confirmed to be SCC or squamous intraepithelial neoplasia. We examined AIRE expression in these esophageal tumors. Although barely detectable in normal epithelium, AIRE expression was distinctively observed in neoplastic cells (Fig 1B, 1C). Nuclear dots as seen in the mTECs were evident in neoplastic cells based on immunofluorescence staining, but they were fewer than in the mTECs (Fig 1A, 1B). In chromogenic detection, cytoplasmic staining was observed in addition to nuclear staining (Fig 1C), the significance of which was uncertain. We examined KRT17 expression in tumor tissues by immunohistochemical staining. KRT17 was barely detected in normal squamous epithelium, but it was clearly observable in tumor tissues, in which it colocalized with AIRE (Fig 1C).

### AIRE is upregulated in human OSCC tissues and cell lines

We first examined AIRE expression in human OSCC using 7 OSCC cell lines and cultured primary keratinocytes as a normal reference. AIRE expression was examined by RT-PCR using an intron-spanning primer set targeting *AIRE*. Gene-specific primers were designed and validated (data not shown). *AIRE* mRNA was detected by RT-PCR in all the OSCC cell lines examined as well as in normal keratinocytes (Fig 2A). Northern blot analysis failed to detect *AIRE* in any cells, suggesting a low amount of transcripts.

**Fig 2.**
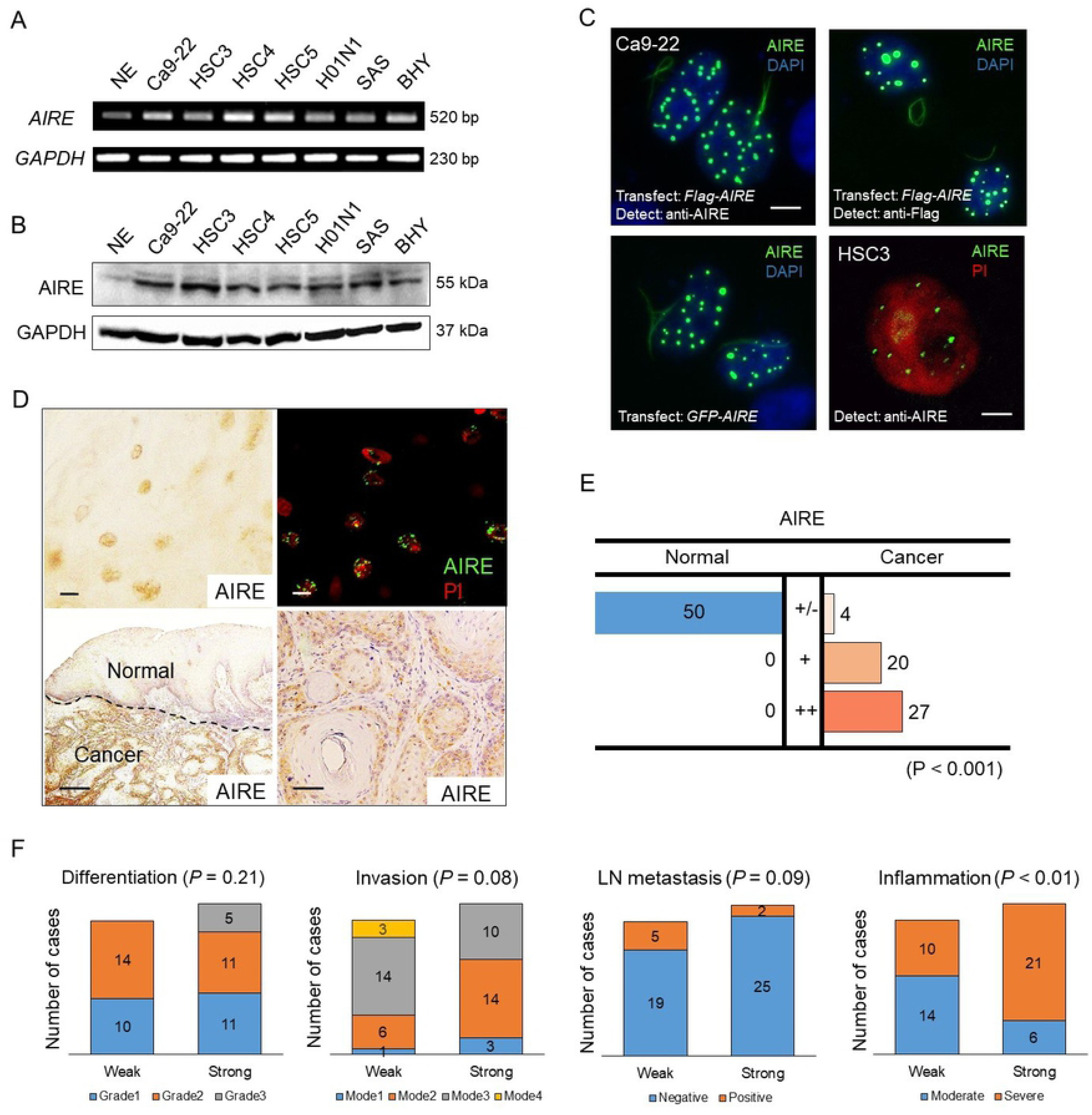
AIRE expression in OSCC. A) RT-PCR analysis of various oral cancer cell lines. NE: normal epithelium from gingiva. GAPDH was used as a positive control. B) Western blot analysis of OSCC lysates using anti-AIRE antibody. GAPDH was used as a loading control. NE: normal epithelial primary cultured cells derived from neonatal foreskin. C) Nuclear localization of AIRE in cultured cells. Upper left and right panels: Ca9-22 cells were transfected with *Flag-AIRE*. The cells were fixed and subjected to immunofluorescence staining. Lower left panel: Ca9-22 cells were transfected with *GFP-AIRE*. The cells were examined under ultraviolet light. Lower right panel: endogenous AIRE in the nucleus of an HSC3 cell as revealed by immunofluorescence staining using laser scanning microscopy. PI: propidium iodide. Scale bar: 10 µm in upper left panel, 5 µm in lower right panel, scale bars were omitted in upper right and lower left panels. D) Representative images of immunohistochemical staining of AIRE in OSCC tissue specimens. Upper left, lower left, lower right panels: chromogenic detection using DAB. Upper right panel: fluorescence detection. Scale bar: 10 µm (upper left, upper right), 500 µm (lower left), 50 µm (lower right). E) Summary of immunohistochemical evaluation of AIRE in OSCC tissue specimens. Expression was compared between normal squamous epithelium and cancer in the same specimen and was scored as +/–: no upregulation in cancer; +: upregulation in cancer; ++: strong upregulation in cancer. Mann-Whitney U test. F) Comparison of AIRE expression with clinicopathological parameters (Mann-Whitney U test). “Weak” refers to cases with the score +/– or + in E), “strong” refers to the cases with score ++ in E).

Western blot analysis using the anti-AIRE antibody revealed that all the OSCC cell lines examined expressed AIRE at substantially higher levels than did normal keratinocytes, which hardly expressed AIRE (Fig 2B). We further investigated the cellular localization of AIRE. For validation of the antibody in cultured-cell staining, 293A or Ca9-22 cells were transfected with *Flag-AIRE* and then stained using anti-AIRE or anti-FLAG antibody 48 h after transfection. Both antibodies produced identical nuclear-dot staining patterns (Fig 2C). The same nuclear localization pattern was observed in live cells expressing GFP-AIRE, and protein distribution was not altered by fixation. Although endogenous AIRE expression was weaker than exogenous expression and the size and number of nuclear dots were lower than those in transfected cells, nuclear dots in HSC3 cells were positive for AIRE as indicated by laser scanning microscopy (Fig 2C). Next, we examined AIRE expression in 50 OSCC tissue specimens. AIRE was detected by immunohistochemical staining in the nuclei and cytoplasm of cancer cells. Immunofluorescence staining also revealed a nuclear dot pattern of AIRE, with substantially less cytoplasmic signal (Fig 2D). Therefore, we regarded the cytoplasmic staining as nonspecific and scored AIRE expression in OSCC based on the nuclear staining intensity in comparison to normal epithelium. Twenty-seven cases showed a distinctive increase (“++”), 20 cases showed a weak increase (“+”), and four cases showed unremarkable change (“+/–”) in AIRE expression. These results indicated that AIRE expression is significantly increased in OSCC compared to normal epithelium (Fig 2E). Next, we examined whether AIRE expression was associated with clinicopathological parameters, including the differentiation grade, invasion pattern, lymph node metastasis, and level of inflammatory cell infiltration in the cancer stroma. OSCC with strong AIRE expression tended to exhibit severer inflammation (Fig 2F).

### AIRE promotes KRT17 expression

As both AIRE and KRT17 were induced in OSCC, we hypothesized that AIRE regulates *KRT17* expression. To investigate the effect of AIRE on *KRT17* expression, we used 293A cells, which lack expression of both. On transfection, AIRE localized mainly to the nuclei, in distinctive dots (Fig 3A). AIRE expression was also confirmed by western blot analysis (Fig 3B). RT-PCR revealed that *KRT17* was induced in *Flag*-*AIRE*-transfected, but not in mock-transfected cells (Fig 3C). Next, we examined the effect of constitutive expression of AIRE by generating stable transformants (293/AIRE+). KRT17 expression was significantly induced in three independent 293/AIRE+ clones as revealed by western blot analysis (Fig 3D).

**Fig 3.**
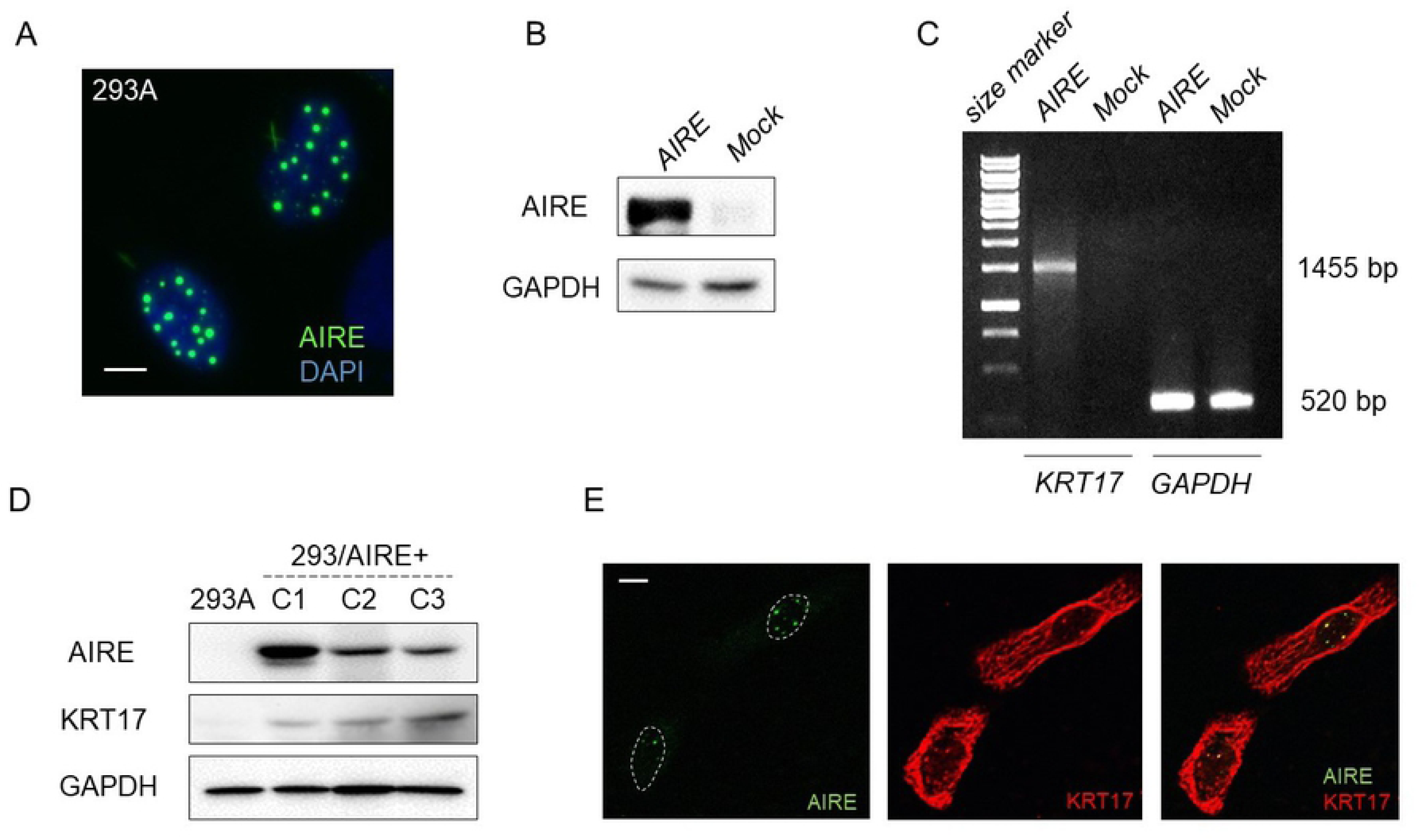
AIRE upregulates KRT17. A) Immunofluorescence examination of 293A cells transfected with *Flag-AIRE*. Scale bar: 5 µm. B) Western blot analysis of 293A cells transiently transfected with *Flag-AIRE*. GAPDH was used as a loading control. C) RT-PCR analysis of KRT17 in 293A cells 72 h after transfection with *Flag-AIRE*. *GAPDH* was used as an internal control. D) Western blot analysis for KRT17 expression in clones C1, C2, C3 of 293A cells stably expressing AIRE using the anti-keratin 17 antibody (EPR1624Y, Abcam). GAPDH was used as a loading control. E) Immunofluorescence double staining of HSC3 cells using the anti-AIRE antibody and a polyclonal anti-keratin 17 antibody (#1900, Epitomics). This anti-keratin 17 antibody was not used in other experiments. Scale bar: 5 µm.

A previous study reported that KRT17 colocalized with AIRE in nuclei of A431 cells (derived from SCC of the skin) treated with leptomycin B, an inhibitor of nuclear export [6]. We checked whether AIRE and KRT17 colocalized in OSCC cells under natural conditions. Four different antibodies against human KRT17, along with the anti-AIRE antibody, were used for immunofluorescence double staining. Three antibodies demonstrated identical filamentous localization of KRT17 in the cytoplasm of HSC3 cells, but no nuclear KRT17. Only one antibody (polyclonal, #1900, Epitomics) revealed a nuclear dot pattern of KRT17 expression, which overlapped with AIRE signal (Fig 3E). However, we found that this polyclonal KRT17 antibody crossreacted with AIRE in western blot analysis, indicating that the nuclear signal was caused by crossreaction of this polyclonal KRT17 antibody with AIRE protein.

### AIRE promotes the expression of STAT1, ICAM1, CXCL10, CXCL11, and MMP9

To investigate the effect of AIRE on other genes, we performed cDNA microarray analysis to screen for genes upregulated by AIRE. Genes whose expression was increased by more than 2-fold in AIRE-expressing compared to control 293A cells in at least three microarray spots are listed in Fig 4A. Among them, we focused on *STAT1* and *ICAM1* because these genes 1) have been detected at considerable levels in our previous cDNA microarray analysis of HSC3 cells [18]; 2) reportedly are overexpressed and play important roles in OSCC [19, 20]; 3) are related to KRT17 in the pathogenesis of psoriasis [21]; and 4) KRT17 has been shown to be upregulated in keratinocytes in a STAT1-dependent manner [22, 23]. Western blot analysis confirmed the upregulation of STAT1 and ICAM1 in AIRE-expressing clones (Fig 4B). Phosphorylated STAT1 increased in proportion to STAT1 expression.

**Fig 4.**
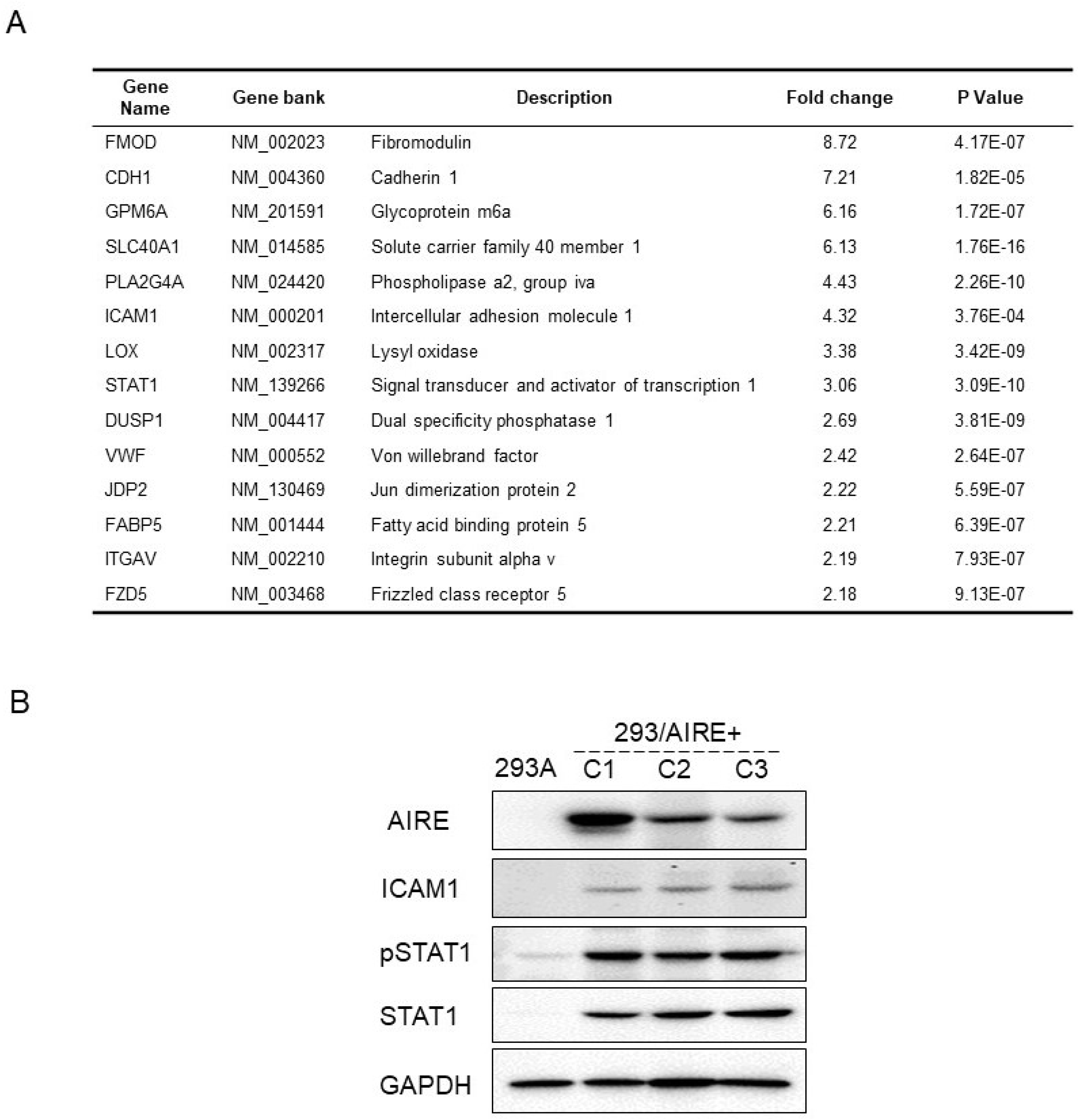
AIRE promotes the expression of ICAM1 and STAT1 in 293A cells. A) Genes that were upregulated more than 2-fold in in 293/AIRE+ (C-3) in comparison to non-transfected 293A cells in at least three spots on the cDNA microarray. B) Western blot analysis of ICAM1 and STAT1 in 293/AIRE+ cells. The results for AIRE and GAPDH are the same as that used in Fig 3B.

To further explore the role of AIRE in OSCC, we disrupted AIRE expression in HSC3 cells. First, we tried transient transfection of siRNA, but the knockdown efficiency at the protein level was less than 20%. Therefore, we used the CRISPR-Cas9 system for gene knockout, and we established three independent *AIRE*-knockout HSC3 cell clones (HSC3/AIRE-C1, C2, and C3, Fig 5A). There was no significant difference in cell proliferation between HSC3/AIRE cells and HSC3 control cells (Fig 5B). A transwell migration assay revealed that the migration rate was significantly suppressed in HSC3/AIRE-C1 and C2 compared to control cells (Fig 5C). HSC3/AIRE-C3 exhibited a lower rate of migration, although the difference was not statistically significant. Western blot analysis revealed that KRT17 expression was decreased in all HSC3/AIRE^−^ clones compared to the control (Fig 5A). ICAM1, STAT1, and phosphorylated STAT1 (pSTAT1) expression were also decreased in HSC3/AIRE compared to control cells.

**Fig 5.**
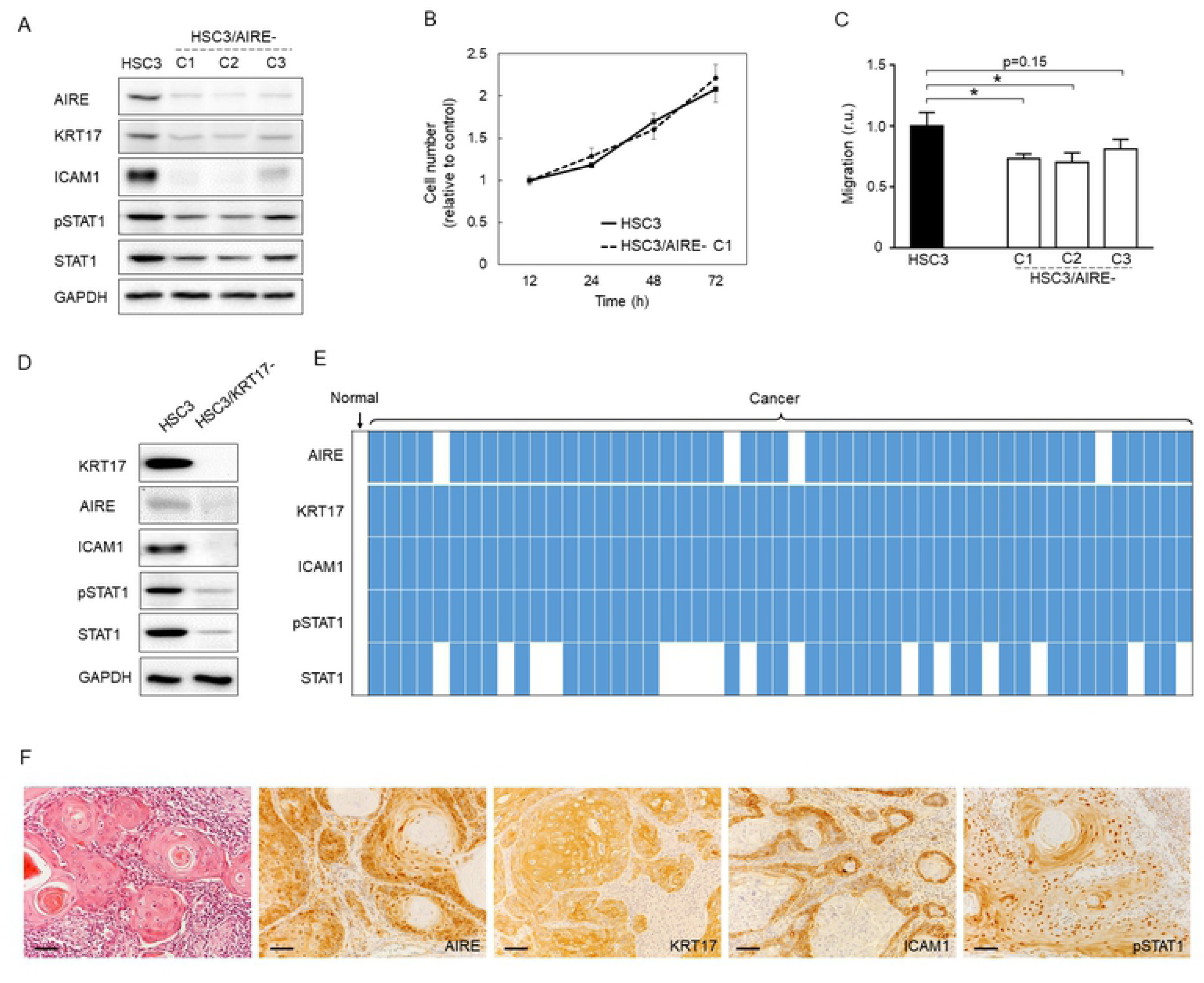
KRT17, STAT1, and ICAM1 are upregulated in OSCC. A) Western blot analysis of KRT17, ICAM1, pSTAT1, and STAT1 in HSC3/AIRE– clones (C1, C2, C3). GAPDH was used as a loading control. B) Cell proliferation of HSC3 and HSC3/AIRE– C1 measured using Cell Counting Kit-8 (Dojindo Laboratories). Data are shown as the mean + SEM of biological triplicates.C2 and C3 also showed similar proliferation rates as HSC3. Student’s *t*-test. C) Transwell migration assay. Data are shown as the mean + SEM of biological triplicates. Student’s *t*-test, * *P* < 0.05. D) Western blot analysis of AIRE, ICAM1, pSTAT1, and STAT1 in HSC3/KRT17– cells. GAPDH was used as a loading control. E) Schematic summary of immunohistochemical expression or AIRE, KRT17, ICAM1, pSTAT1, and STAT1 in 50 cases of OSCC. Horizontal grids represent the cases. Filled squares denote upregulation compared to normal epithelium in the same specimen. Blank squares denote no upregulation, i.e., similar staining intensity as in normal epithelium. F) Representative images of immunohistochemical staining. Scale bar: 50 µm.

The relationship between AIRE and KRT17 was further explored using previously established KRT17-knockout HSC3 cells (HSC3/KRT17^−^) [24]. In HSC3/KRT17^−^ cells, the expression of AIRE, ICAM1, and STAT1 was remarkably reduced compared to control cells (Fig 5D). To confirm these observations, we investigated the expression of KRT17, ICAM1, and pSTAT1 in 50 OSCC tissue specimens by immunohistochemistry. In most cases, KRT17, ICAM1, and pSTAT1 expression was increased in OSCC tissue compared to normal epithelium (Fig 5E). These findings supported the results of cell-culture experiments showing that AIRE promoted KRT17, STAT1, and ICAM1 expression. AIRE and KRT17 cooperatively stimulate the expression of the cancer-related proinflammatory genes *MMP9*, *CXCL10*, and *CXCL11* [6]. In line herewith, we found significantly reduced expression of *MMP9*, *CXCL10*, and *CXCL11* in HSC3/KRT17^−^ (Fig 6A) and HSC3/AIRE^−^ cells (Fig 6B), and significant upregulation of *MMP9*, *CXCL10*, and *CXCL11* in AIRE-expressing 293A cells (Fig 6C). These results suggested a common mechanism that regulates the expression of these cancer-related genes.

**Fig 6.**
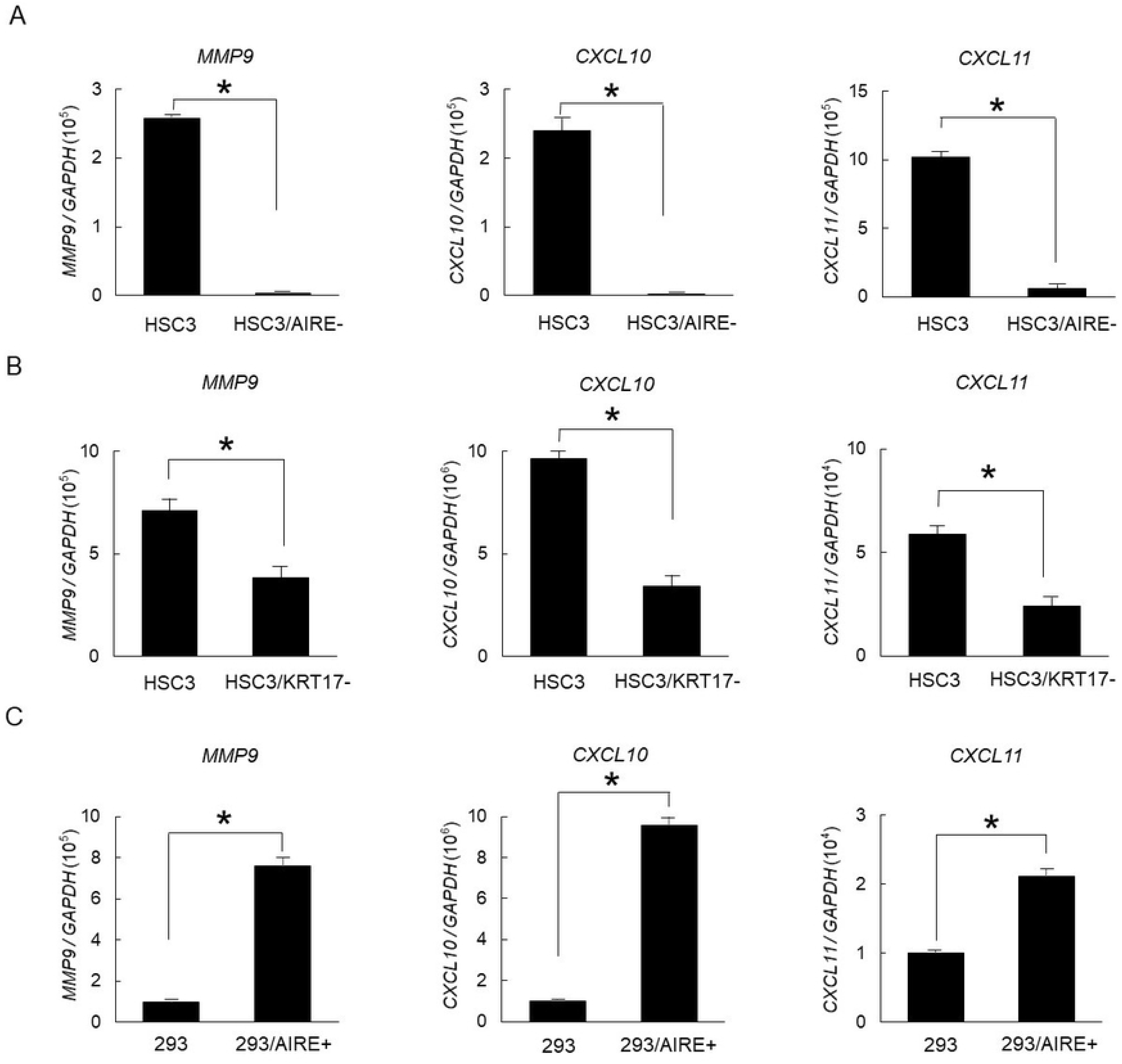
Relative expression of proinflammatory genes. Gene expression in A) HSC3/AIRE–, B) HSC3/KRT17–, and C) 293/AIRE+ in comparison to respective non-transformed cells as measured by real-time RT-PCR. *GAPDH* was used as an internal control. Data are shown as the mean + SEM of technical triplicates and are representative of three independent experiments. Student’s *t*-test. \**P* < 0.05.

### AIRE promotes cancer-related gene expression in cooperation with ETS1

Nuclear dots containing AIRE reportedly are different from nuclear dots containing Sp100, the closest homologue of AIRE. However, we found that AIRE and Sp100 colocalized in nuclear dots as revealed by double staining for AIRE and Sp100 in *Flag*-*AIRE*-transfected cells (S1 Fig). Sp100 physically interacts with ETS1 and mediates its transcriptional activity [25, 26]. ETS1 is a member of the ETS family of transcription factors that physically and functionally interact with numerous molecules to control transcription [27]. Furthermore, various genes that we found to be positively correlated with AIRE in terms of expression (i.e., *KRT17*, *STAT1*, *ICAM1*, *MMP9*, *CXCL10*, and *CXCL11*) have been previously reported as remarkably upregulated in cDNA microarray analysis of epidermal keratinocytes from *Ets1* transgenic mice [28]. Therefore, we hypothesized that AIRE and ETS1 cooperatively contribute to expression regulation of these genes. The OSCC lines expressed ETS1, along with STAT1 and ICAM1, at various levels (Fig 7A). We transiently transfected *Flag*-*AIRE* or/and *Ets1* into 293A cells, which do not express ETS1 at a detectable level as indicated by western blot analysis, and we examined the expression of the aforementioned genes. Western blot analysis revealed increases in STAT1 and ICAM1 expression upon overexpression of AIRE or/and ETS1 (Fig 7B, 7C). Coexpression of AIRE and ETS1 resulted in enhanced expression of STAT1. Quantitative RT-PCR revealed that *CXCL10* and *MMP9* expression was upregulated by ETS1 expression, and even more so upon cotransfection with *AIRE* (Fig 7D).

**Fig 7.**
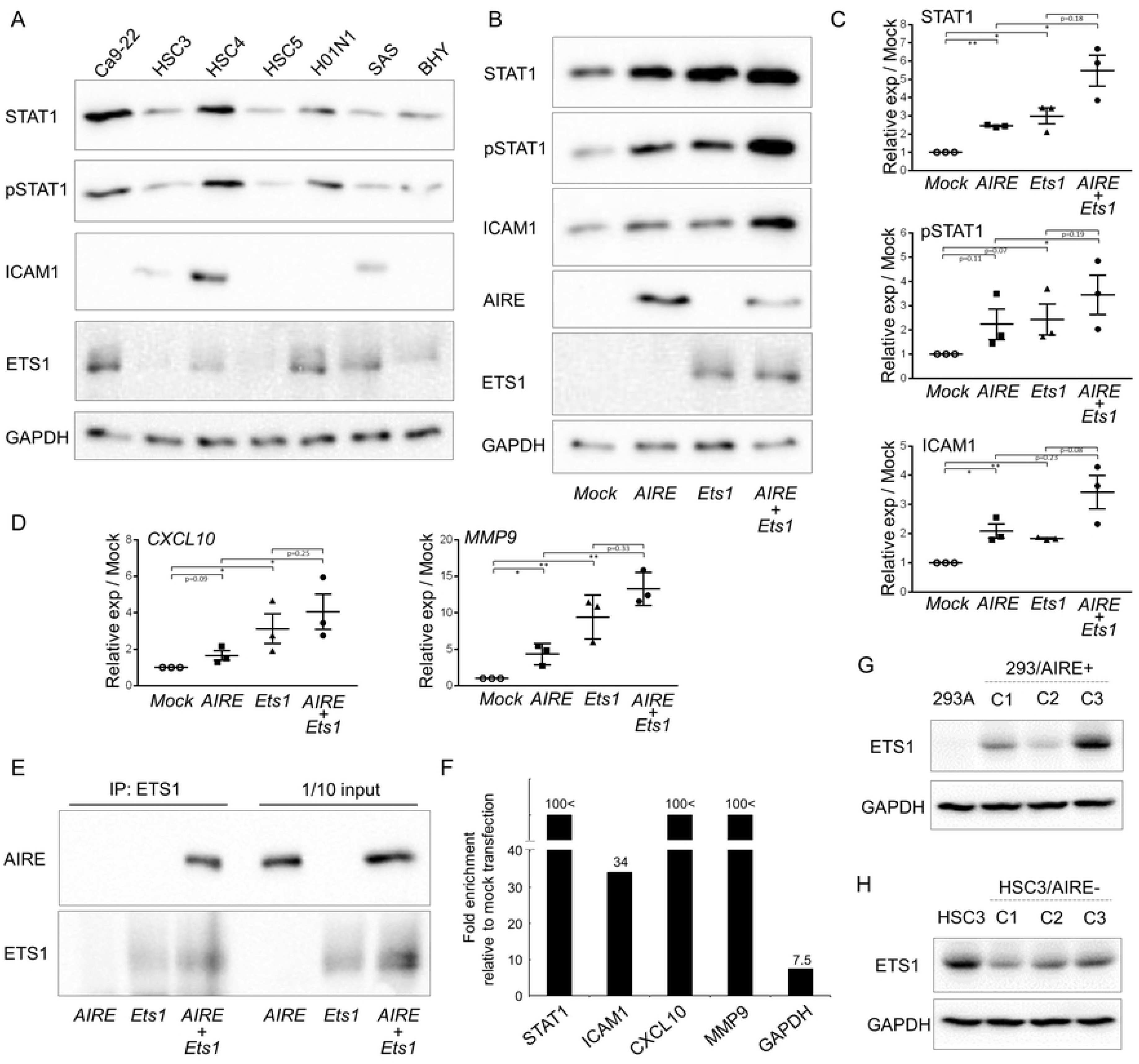
Physical and functional interaction of AIRE and ETS1. A) Expression of STAT1, pSTAT1, ICAM1, and ETS1 in various OSCC cell lines as revealed by western blot analysis. GAPDH was used as a loading control. B) Expression of STAT1, pSTAT1, and ICAM1 in 293A cells 48 h after transfection with *AIRE* or/and *ETS1*, or mock transfection. The blot shown is representative of three independent experiments. C) Densitometrical analysis of data in B). Data are shown as mean ± SEM of technical triplicates and are representative of three independent experiments. Ratio paired t-test. * *P* < 0.05; ** *P* < 0.01. D) Relative gene expression of CXCL10 and MMP9 in comparison to mock transfection as revealed by real time PCR. Data are shown as mean ± SEM of triplicate wells and are representative of two independent experiments. Ratio paired t-test. * *P* < 0.05; ** *P* < 0.01. E) Immunoprecipitation and western blot analysis of ETS1 and AIRE. Formaldehyde crosslinking was performed prior to lysis. The lysates were sonicated to shear the DNA. Immunoprecipitation with the anti-ETS1 antibody was performed, and recovered protein was examined by western blot analysis. One tenth of the sample used for immunoprecipitation was loaded as a control. F) ChIP assay of 293A cells transiently transfected with AIRE. Enrichment of promoter fragments was measured by real-time PCR. Values were standardized to the input, and then to mock transfection. Relative values higher than 100 are indicated as 100< and a scale break was used in the Y-axis to allow intuitive interpretation of the relative expression. GAPDH was used as a reference. Data are representative of three independent experiments. G, H) ETS1 expression in G) 293/AIRE+ and H) HSC3/AIRE– as revealed by western blot analysis. GAPDH was used as a loading control.

To examine the physical interaction between AIRE and ETS1, we cotransfected *Flag-AIRE* or/and *Ets1* into 293A cells and used these cells for immunoprecipitation. Conventional immunoprecipitation using anti-ETS1 antibody did not show coprecipitation of AIRE with ETS1 (data not shown), which indicates that their interaction is weak or their functional cooperation is not mediated through direct physical interaction. ETS1 and AIRE may be recruited to nearby chromatin regions. To check this possibility, we performed a modified immunoprecipitation assay. Formaldehyde crosslinking was performed before cell lysis, and the DNA was sheared by sonication. Immunoprecipitation was conducted using the anti-ETS1 antibody, and recovered protein was examined by western blot analysis. AIRE coprecipitated with ETS1 (Fig 7E). We next performed conventional ChIP assays to assess whether AIRE was recruited to the promoters of the aforementioned genes. ChIP-quantitative PCR revealed that promoter fragments of *STAT1*, *ICAM1*, *CXCL10*, and *MMP9* were enriched in the AIRE precipitate (Fig 7F). Although transient expression of *AIRE* did not lead to noticeable upregulation of ETS1 in 293A cells (Fig 7B), ETS1 was induced in stable 293/AIRE+ clones (Fig 7G). In contrast, ETS1 expression was decreased in HSC3/AIRE^−^ cells (Fig 7H).

To further examine the association between ETS1 and AIRE, we evaluated their cellular localization. In Ca9-22 cells, ETS1 was observed diffusely in the nucleus (Fig 8A), which partially overlapped with transfected AIRE (Fig 8A). We next examined the effect of ETS1 on the cellular distribution of AIRE in 293A cells. Following *AIRE* transfection, the cellular distribution of AIRE changed over time. At 12 h after transfection, most AIRE was present in the cytoplasm, reflecting initial protein synthesis from the plasmid (Fig 8B). At 24 h after transfection, nuclear localization of AIRE was observed in a small number of cells, whereas most AIRE protein was still observed in the cytoplasm. At 48 h after transfection, AIRE was predominantly observed as dots in the nucleus (Fig 8B), indicating that protein synthesis from the transfected plasmid had ceased and all AIRE protein had translocated from the cytoplasm to the nucleus. When *Ets1* was cotransfected, AIRE distribution was different from that in cells transfected with AIRE alone; at 36 h, most AIRE protein was observed in the nuclei of cells expressing ETS1, in contrast to the largely cytoplasmic localization in *AIRE*-single transfectants. We evaluated the cellular localization of AIRE in 100 cells 36 h after transfection and we classified individual cells into three classes—nuclear dominant, mixed, and cytoplasmic dominant—according to the cellular distribution of AIRE (Fig 8C). *AIRE*/*Ets1* cotransfection resulted in more cells with nuclear AIRE than *AIRE*-single transfection. Taken together, these results implicates that AIRE was recruited to the chromosomal regions where ETS1 initiated transcription.

**Fig 8.**
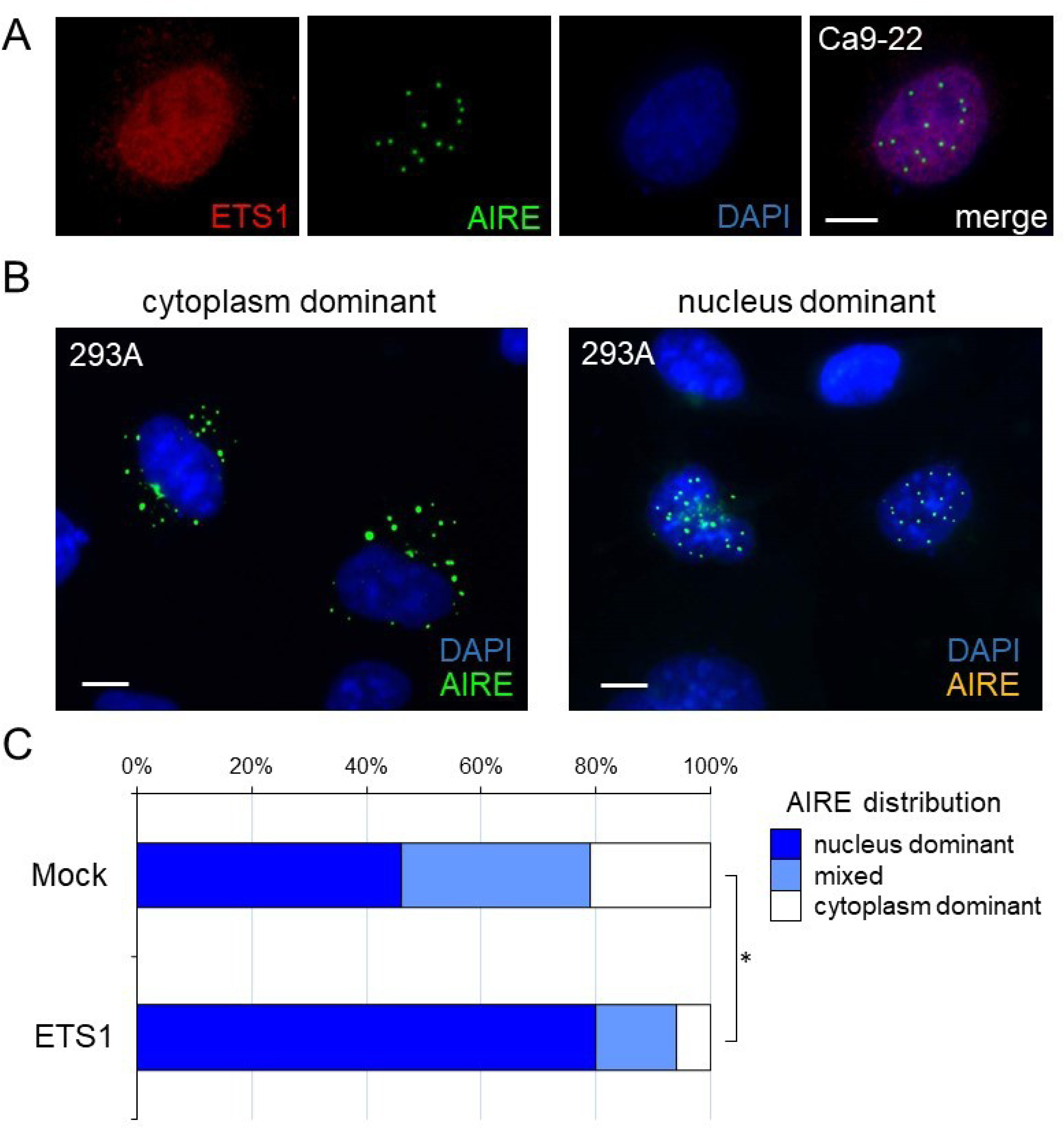
Nuclear translocation of AIRE in the presence of ETS1. A) Nuclear distribution of ETS1 (red) and AIRE (green). Ca9-22 cells were transfected with *GFP-AIRE*. Immunofluorescence staining using the anti-ETS1 antibody was performed 48 h after transfection. Nuclei were counterstained with DAPI. Scale bar: 5 µm. B) Representative patterns of cellular distribution of AIRE. 293A cells were transfected with *GFP-AIRE*. Scale bar: 5 µm. C) Cellular distribution of AIRE in 293A cells 36 h after transfection with *GFP-AIRE* together with empty vector or *Ets1*. The cellular distribution of AIRE was evaluated 36 h after transfection as three patterns; nuclear dominant, mixed, and cytoplasmic dominant One hundred randomly chosen cells were evaluated. Mann-Whitney U test. * *P* < 0.05. Data are representative of three independent experiments.

## Discussion

Although accumulating evidence suggests that AIRE is expressed in extrathymic tissues, there are remarkable discrepancies across studies, depending on the techniques used. *AIRE* mRNA has been detected by RT-PCR in lymph nodes, tonsils, gut-associated lymphoid tissues, spleen, liver, testes, and ovaries [29]. Thymic *AIRE* mRNA expression is the highest and could also be detected by northern blotting [30, 31]. As studies have concentrated mainly on lymphoid tissues, the skin and upper digestive tract mucosa have been left unexplored. Only a few studies have reported AIRE expression in epithelial tissues. *In-situ* hybridization and immunohistochemistry failed to detect AIRE expression in human skin [8], whereas *AIRE* expression has been detected by RT-PCR in HaCaT keratinocytes [11]. Hobbs and colleagues demonstrated *AIRE* mRNA in A431 cells treated with 12-O-tetradecanoylphorbol 13-acetate (TPA) by RT-PCR. TPA is an activator of the protein kinase C pathway, which induces inflammation, and acts as a strong carcinogen when topically applied to skin. *Aire* mRNA expression was barely detectable in normal mouse skin by *in-situ* hybridization, but was induced in the epidermis of TPA-treated mouse ear and genetically induced mouse skin tumor. Still, induced epidermal *Aire* expression was substantially lower than that in the thymus [6]. Our results showing that AIRE is expressed in OSCC are consistent with these previous observations, and suggest that AIRE is induced in the epithelium under pathological conditions.

We examined the expression of several tissue-specific antigens, including PTH, INS, GH1, which are induced in mTECs in an AIRE-dependent manner [7], in OSCC cell lines. None of these were detected in the OSCC cell lines examined or induced in AIRE-overexpressing cells (unpublished results). This is plausible considering the presumptive function of AIRE in reinstigating suspended transcription. Instead, we found evidence of upregulation of KRT17, STAT1, and ICAM1 by AIRE. The coordinated overexpression of KRT17, STAT1, and ICAM1 in OSCC is consistent with findings in previous studies [20,24,29,32,33]. Close correlations between KRT17, STAT1, and ICAM1 expression in epithelial cells have been demonstrated in previous studies. Interferon gamma activates STAT1 and induces KRT17 expression in keratinocytes, which has been considered part of a crucial pathogenic mechanism for psoriasis [22, 23]. In addition, interferon gamma induces ICAM1 expression through activation of STAT1 [34–36]. Although the regulatory mechanism of STAT1 transcription is unclear, our finding that AIRE induces STAT1 expression suggests that AIRE may increase the expression of KRT17 and ICAM1 through STAT1 upregulation.

We found that AIRE stimulates *MMP9*, *CXCL10*, and *CXCL11*, which is consistent with findings by Hobbs and colleagues [6]. These authors demonstrated that AIRE and KRT17 are recruited to a specific region in the promoters of these proinflammatory genes carrying an NF-kappaB consensus motif [6]. We failed to confirm a direct physical interaction between AIRE and KRT17 in our experimental system. Thus, KRT17-dependent transcriptional control by AIRE may be a context-dependent phenomenon.

Interestingly, the same spectrum of genes whose expression was dependent on AIRE in keratinocytes under stress in our study, including *Krt17*, *Stat1*, *Icam1*, *Mmp9*, *Cxcl10*, and *Cxcl11*, was reportedly induced in keratinocytes of *Ets1* transgenic mice [28]. Microarray analysis of *Ets1* transgenic in comparison with wild-type mouse epidermis revealed the upregulation of *Krt17* (2.9-fold), *Stat1* (2.4-fold), *Icam1* (2.8-fold), *Mmp9* (8.9-fold), *Cxcl10* (8.2-fold), and *Cxcl11* (3.3-fold) in the former [28]. Unfortunately, data on *Aire* expression were lacking in this previous microarray study, but the results in this and previous studies suggest that AIRE belongs to this group of genes that are induced in epithelial cells under stress condition.

AIRE regulates the transcription of numerous divergent genes whose expression is driven by various promoters and transcription factors. This suggests that AIRE is a broad transcriptional activator, rather than a specific DNA-binding transcription factor. The mechanism underlying this promiscuous transcriptional activation is not fully understood, but it has been proposed that AIRE is recruited to RNA polymerase II-rich regions where transcription of AIRE-responsive genes is initiated but interrupted, and AIRE releases the stalled RNA polymerase II complex to resume transcription [37–39]. Further, AIRE has been suggested to be preferentially recruited to so-called super-enhancers; genomic regions rich in enhancers and transcriptional regulators [40]. Through its global transcriptional activation, AIRE might assist cancer-associated gene expression and thus promote the cancer phenotype.

The regulatory mechanism of AIRE expression has not been clearly understood. The *AIRE* minimal promoter region contains binding domains for Sp1, AP-1, NF-Y [41], and Ets transcription factors [42]. Indeed, ETS1 and ETS2 enhanced TPA-induced *AIRE* promoter activity in HeLa cells [42], although transient overexpression of ETS1 alone did not induce AIRE in 293A cells. Further research is needed to elucidate the details of the mechanism of AIRE induction in cancer cells.

In conclusion, we demonstrated a novel role of AIRE in extra-thymic tissue under pathologic condition. AIRE is upregulated in OSCC and promotes the expression of cancer-related genes, at least in part by functional interaction with ETS1.

## Acknowledgments

This work was supported by JSPS KAKENHI (grant Nos. JP25462848 and JP16K11438) and in part by a grant for Private University Branding Project to Tokyo Dental College from MEXT.

